# On the Colijn-Plazzotta numbering scheme for unlabeled binary rooted trees

**DOI:** 10.1101/2020.06.16.155184

**Authors:** Noah A. Rosenberg

## Abstract

Colijn & Plazzotta (*Syst. Biol.* 67:113-126, 2018) introduced a scheme for bijectively associating the unlabeled binary rooted trees with the positive integers. First, the rank 1 is associated with the 1-leaf tree. Proceeding recursively, ordered pair (*k*_1_, *k*_2_), *k*_1_ ⩾ *k*_2_ ⩾ 1, is then associated with the tree whose left subtree has rank *k*_1_ and whose right subtree has rank *k*_2_. Following dictionary order on ordered pairs, the tree whose left and right subtrees have the ordered pair of ranks (*k*_1_, *k*_2_) is assigned rank *k*_1_(*k*_1_ − 1)*/*2 + 1 + *k*_2_. With this ranking, given a number of leaves *n*, we determine recursions for *a*_*n*_, the smallest rank assigned to some tree with *n* leaves, and *b*_*n*_, the largest rank assigned to some tree with *n* leaves. For *n* equal to a power of 2, the value of *a*_*n*_ is seen to increase exponentially with 2*α*^*n*^ for a constant *α* ≈ 1.24602; more generally, we show it is bounded *a*_*n*_ < 1.5^*n*^. The value of *b*_*n*_ is seen to increase with 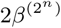 for a constant *β* ≈ 1.05653. The great difference in the rates of increase for *a*_*n*_ and *b*_*n*_ indicates that as the index *v* is incremented, the number of leaves for the tree associated with rank *v* quickly traverses a wide range of values. We interpret the results in relation to applications in evolutionary biology.

**Mathematics subject classification:** 05C05, 92B10, 92D15

## 1 Introduction

For a given number of leaves *n* ⩾ 2, the unlabeled binary rooted trees with *n* leaves can be obtained recursively (Table 1). For fixed *n*, we enumerate all possible pairings of a subtree of size *k* leaves with a subtree of size *n* − *k* leaves, for each *k* from 1 to 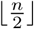. For each 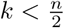., each pairing of a subtree of size *k* and a subtree of size *n* − *k* generates a distinct unlabeled binary rooted tree; for even *n* and 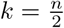., we enumerate pairings of distinct subtrees of size 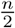. and pairings of identical subtrees of size 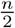.

**Table 1.**
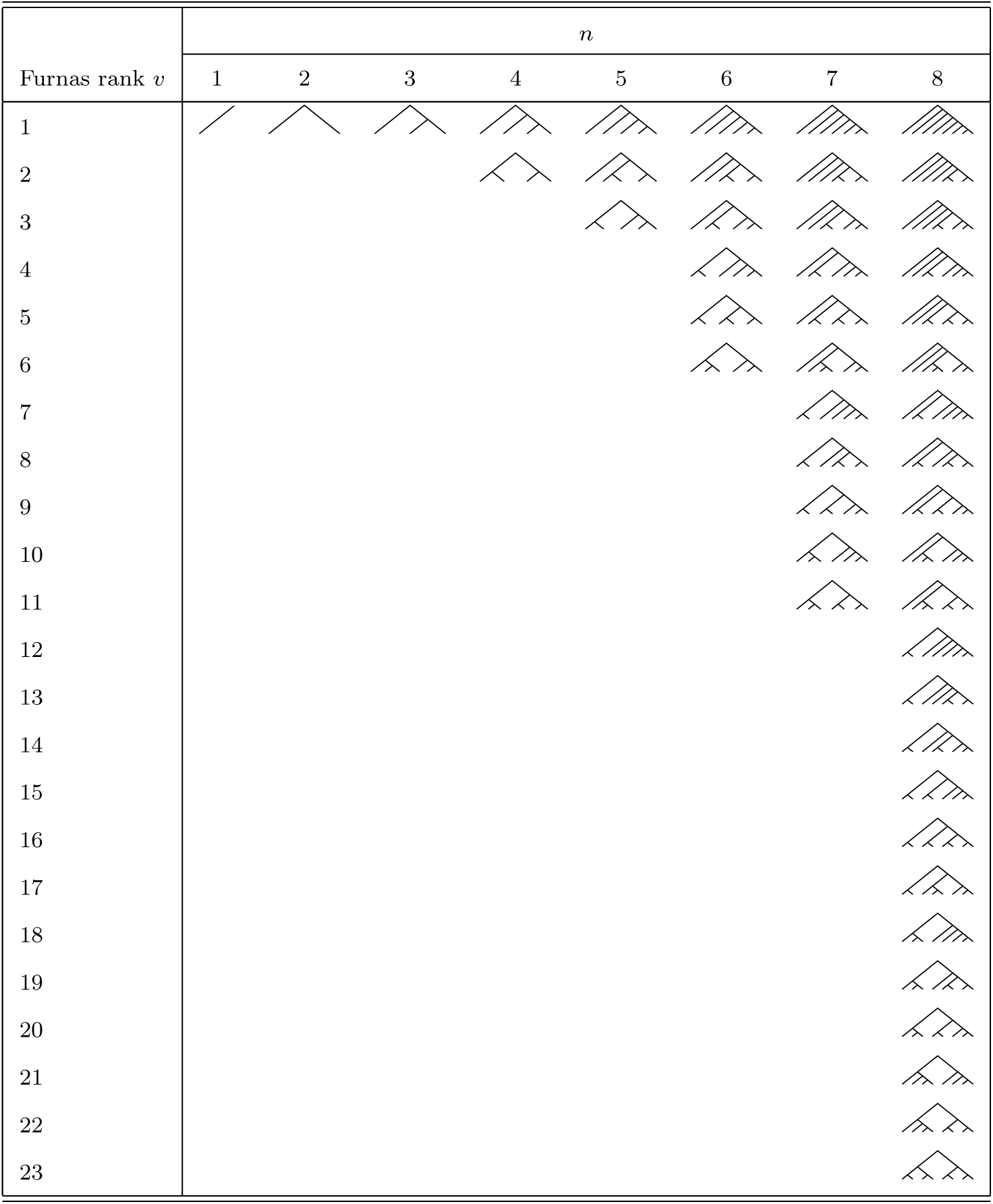
Furnas ranks of unlabeled binary rooted trees with 1 ⩽ *n* ⩽ 8 leaves.

Letting *U*_*n*_ denote the number of unlabeled binary rooted trees with *n* leaves, we have [10, p. 29]

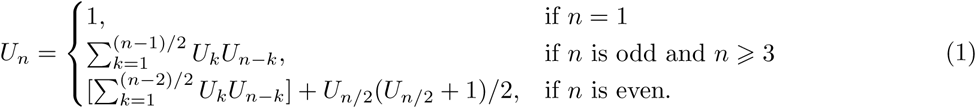

The sequence of values *U*_*n*_, the Wedderburn-Etherington numbers, begins from *n* = 1 with 1, 1, 1, 2, 3, 6, 11, 23, 46, 98, 207, 451, 983, 2179, 4850 (Table 2, A001194 in OEIS). *U*_*n*_ is straightforward to calculate from *U*_1_, *U*_2_, …, *U*_*n*−1_ via the recursion in eq. 1. However, no closed-form expression is known.

**Table 2.**
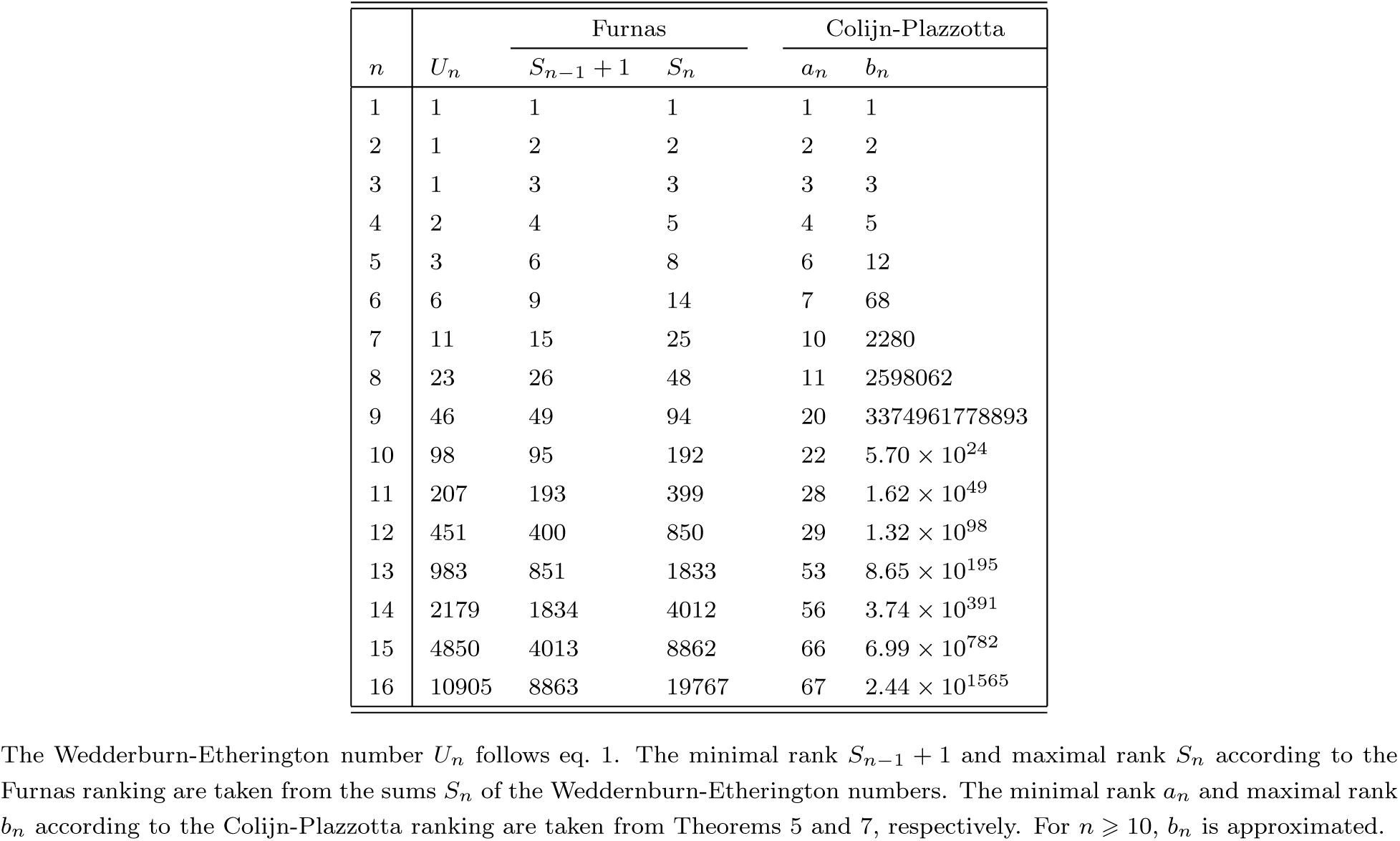
Minimal and maximal Furnas and Colijn-Plazzotta ranks among unlabeled binary ranked trees with 1 ⩽ *n* ⩽ 16 leaves.

For a fixed value of *n*, the unlabeled binary rooted trees can be enumerated in the sequence in which they appear in the recursion. According to the ranking scheme of Furnas [11] for trees of size *n* leaves, 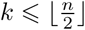. is viewed as the size of the left subtree of a tree of size *n* ⩾ 2 and *n* − *k* is the size of the right subtree. Trees with *n* leaves that have a lower value of *k* are assigned lower rank. Trees with *n* leaves that have the same value of *k* are ordered by the rank of their left subtree, and trees with *n* leaves that have the same value of *k* and the same left subtree are ordered by the rank of their right subtree. For trees with two distinct subtrees of size 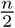, the one with lower Furnas rank appears on the left (Table 1).

The Furnas ranking bijectively associates the unlabeled binary rooted trees with the positive integers. For *n* ⩾ 1, we let 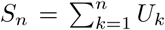. denote the sum of the Wedderburn-Etherington numbers, with *S*_0_ = 0 (A173282 in OEIS). In the bijection, the tree of size *n* with Furnas rank *v*, 1 ⩽ *v* ⩽ *U*_*n*_, is associated with the integer *S*_*n*−1_ + *v*. The trees of size *n* are associated with the integers in [*S*_*n*−1_ + 1, *S*_*n*_] (Table 2).

This bijection based on the Furnas ranking is convenient as a scheme for indexing trees, but the unavailability of a closed form for *U*_*n*_ and hence for *S*_*n*_ makes it difficult to quickly discern the tree associated with a given integer and vice versa. An alternative scheme of Colijn & Plazzotta [4], which also bijectively associates the unlabeled binary rooted trees with the positive integers, addresses this problem.

In the Colijn-Plazzotta ranking, the 1-leaf tree is given rank 1. For *n* ⩾ 2 leaves, the ordered pair (*k*_1_, *k*_2_), *k*_1_ ⩾*k*_2_ ⩾1, is associated with the tree whose left subtree has Colijn-Plazzotta rank *k*_1_ and whose right subtree has rank *k*_2_. Following the dictionary order on ordered pairs, the tree associated with ordered pair (*k*_1_, *k*_2_) is assigned rank *k*_1_(*k*_1_ −1)*/*2+1+*k*_2_. Thus, the Colijn-Plazzotta rank of a tree is obtained recursively from the ranks of its left and right subtrees, and the tree associated with a rank *v* is obtained by identifying the largest *k*_1_ such that *k*_1_(*k*_1_ − 1)*/*2 + 1 < *v* and assigning to rank *v* the tree whose left subtree has rank *k*_1_ and whose right subtree has rank *v* − *k*_1_(*k*_1_ − 1)*/*2 − 1 (Table 3). Note that the left–right orientation of an unlabeled binary rooted tree generally differs for the Furnas and Colijn-Plazzotta rankings.

**Table 3.**
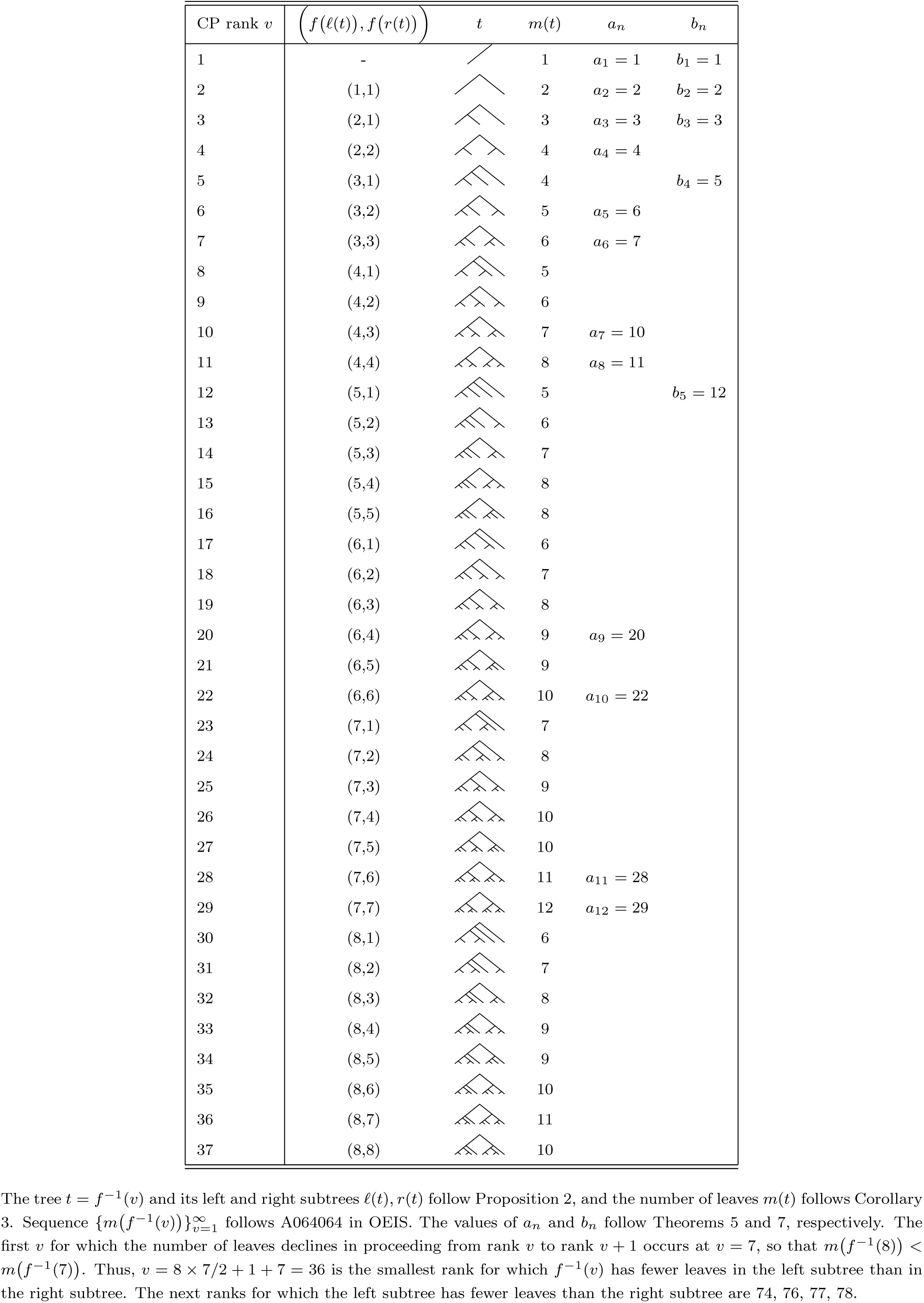
Colijn-Plazzotta ranks of unlabeled binary ranked trees with CP rank 1 ⩽ *v* ⩽ 37.

| CP rank $v$ | $(f(\ell(t)), f(r(t)))$ | $t$ | $m(t)$ | $a_n$ | $b_n$ |
| --- | --- | --- | --- | --- | --- |
| 1 | - | | 1 | $a_1 = 1$ | $b_1 = 1$ |
| 2 | (1,1) | | 2 | $a_2 = 2$ | $b_2 = 2$ |
| 3 | (2,1) | | 3 | $a_3 = 3$ | $b_3 = 3$ |
| 4 | (2,2) | | 4 | $a_4 = 4$ | |
| 5 | (3,1) | | 4 | | $b_4 = 5$ |
| 6 | (3,2) | | 5 | $a_5 = 6$ | |
| 7 | (3,3) | | 6 | $a_6 = 7$ | |
| 8 | (4,1) |  | 5 |  |  |
| 9 | (4,2) |  | 6 |  |  |
| 10 | (4,3) | | 7 | $a_7 = 10$ | |
| 11 | (4,4) | | 8 | $a_8 = 11$ | |
| 12 | (5,1) | | 5 | | $b_5 = 12$ |
| 13 | (5,2) |  | 6 |  |  |
| 14 | (5,3) |  | 7 |  |  |
| 15 | (5,4) |  | 8 |  |  |
| 16 | (5,5) |  | 8 |  |  |
| 17 | (6,1) |  | 6 |  |  |
| 18 | (6,2) |  | 7 |  |  |
| 19 | (6,3) |  | 8 |  |  |
| 20 | (6,4) | | 9 | $a_9 = 20$ | |
| 21 | (6,5) |  | 9 |  |  |
| 22 | (6,6) | | 10 | $a_{10} = 22$ | |
| 23 | (7,1) |  | 7 |  |  |
| 24 | (7,2) |  | 8 |  |  |
| 25 | (7,3) |  | 9 |  |  |
| 26 | (7,4) |  | 10 |  |  |
| 27 | (7,5) |  | 10 |  |  |
| 28 | (7,6) | | 11 | $a_{11} = 28$ | |
| 29 | (7,7) | | 12 | $a_{12} = 29$ | |
| 30 | (8,1) |  | 6 |  |  |
| 31 | (8,2) |  | 7 |  |  |
| 32 | (8,3) |  | 8 |  |  |
| 33 | (8,4) |  | 9 |  |  |
| 34 | (8,5) |  | 9 |  |  |
| 35 | (8,6) |  | 10 |  |  |
| 36 | (8,7) |  | 11 |  |  |
| 37 | (8,8) |  | 10 |  |  |

The tree $t = f^{-1}(v)$ and its left and right subtrees $\ell(t), r(t)$ follow Proposition 2, and the number of leaves $m(t)$ follows Corollary 3. Sequence $\{m(f^{-1}(v))\}_{v=1}^{\infty}$ follows A064064 in OEIS. The values of $a_n$ and $b_n$ follow Theorems 5 and 7, respectively. The first $v$ for which the number of leaves declines in proceeding from rank $v$ to rank $v+1$ occurs at $v=7$ , so that $m(f^{-1}(8)) < m(f^{-1}(7))$ . Thus, $v = 8 \times 7/2 + 1 + 7 = 36$ is the smallest rank for which $f^{-1}(v)$ has fewer leaves in the left subtree than in the right subtree. The next ranks for which the left subtree has fewer leaves than the right subtree are 74, 76, 77, 78.

Here, we study mathematical properties of the Colijn-Plazzotta ranking of the unlabeled binary rooted trees. For fixed *n*, we obtain recursions for the smallest rank *a*_*n*_ assigned to some tree with *n* leaves as well as the largest rank *b*_*n*_. We then study asymptotic properties of *a*_*n*_ and *b*_*n*_.

## 2 The Colijn-Plazzotta ranking

We define the Colijn-Plazzotta ranking more formally. Let *T*_*n*_ be the set of unlabeled binary rooted trees with *n* leaves, and let 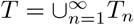 be the set of all unlabeled binary rooted trees. All trees considered here are unlabeled binary rooted trees, and we refer to them simply as *trees*. For a tree *t* ∈ *T*, we let *m*(*t*) denote its number of leaves. For *m*(*t*) ⩾ 2, we let *ℓ*(*t*) and *r*(*t*) denote the left and right subtrees of *t*.

### Definition 1.

*The* Colijn-Plazzotta ranking *for trees t* ∈ *T is a function f* : *T* → ℤ^+^ *that satisfies*

a. *f* (*t*) = 1 *if m*(*t*) = 1, *and*
b. *f* (*t*) = *f* (*ℓ*(*t*))[*f* (*ℓ*(*t*)) − 1]*/*2 + 1 + *f* (*r*(*t*)) *if m*(*t*) ⩾ 2.

We abbreviate the Colijn-Plazzotta ranking as the *CP ranking*. To determine the CP rank of a tree *t*, we require *t* to be written in a canonical form in which *f (ℓ*(*t*)) ⩾ *f (r*(*t*)). In this canonical form, the number of leaves in the left subtree, *m (ℓ*(*t*)), can be greater than, less than, or equal to *m (r*(*t*)) (Table 3). The 1-leaf tree has CP rank 1, and hence, if it is a subtree of the root of *t* and *m*(*t*) ⩾ 3, then it is necessarily the right subtree (for *m*(*t*) = 2, both subtrees have 1 leaf). The 2-leaf tree has CP rank 2, and if it is a subtree of the root of *t* and *m*(*t*) ⩾ 5, then it is the right subtree.

The dictionary order used in the CP ranking has the implication that for two trees *t*_1_, *t*_2_ in canonical form with *f (ℓ*(*t*_1_)) < *f (ℓ*(*t*_2_)), *f* (*t*_1_) < *f* (*t*_2_). For two trees *t*_1_, *t*_2_ in canonical form with *f (ℓ*(*t*_1_)) = *f (ℓ*(*t*_2_)) and *f (r*(*t*_1_)) < *f (r*(*t*_2_)), *f* (*t*_1_) < *f* (*t*_2_).

The CP ranking *f* gives a bijective map between trees and positive integers [4]. Briefly, for injectivity, two distinct trees *t*_1_, *t*_2_ differ in their pair of subtrees, (*ℓ*(*t*_1_)), *(r*(*t*_1_)) ≠ (*ℓ*(*t*_2_)), *r*(*t*_2_)), giving rise to distinct values of *f, f* (*t*_1_) ≠ *f* (*t*_2_). For surjectivity, each positive integer *v* ⩾ 2 has a unique representation in the form *k*_1_(*k*_1_ − 1)*/*2 + 1 + *k*_2_, with *k*_1_, *k*_2_ positive integers and *k*_1_ ⩾ *k*_2_, so that the tree whose subtrees have CP ranks *k*_1_, *k*_2_ is assigned to CP rank *v*.

Given a positive integer *v* ⩾ 2, we identify the tree with CP rank *v* as the tree *t* ∈ *T* whose left subtree is the tree with CP rank *k*_1_(*v*), where *k*_1_(*v*) is the largest integer satisfying *k*_1_(*k*_1_ − 1)*/*2 + 1 < *v*, and whose right subtree is the tree with CP rank *k*_2_(*v*) = *v* − *k*_1_(*v*)[*k*_1_(*v*) − 1]*/*2 − 1. We solve the inequality for *k*_1_.

### Proposition 2.

*The function f* ^−1^ : ℤ^+^ → *T that gives the tree with specified CP rank satisfies*

a. *f* ^−1^(1) *is the tree with one leaf, and*
b. *for v* ⩾ 2, *f* ^−1^(*v*) *is the tree whose left subtree has CP rank* 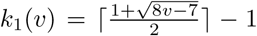 *and whose right subtree has CP rank k*_2_(*v*) = *v* − *k*_1_(*v*)[*k*_1_(*v*) − 1]*/*2 − 1.

Using the function *f* ^−1^ that gives the tree associated with CP rank *v*, we obtain a recursion for the number of leaves possessed by the tree of CP rank *v*.

### Corollary 3.

*The function m* : ℤ^+^ → ℤ^+^ *that gives the number of leaves in the tree with specified CP rank satisfies*

a. *m*(*f* ^−1^(1)) = 1, *and*
b. *for* 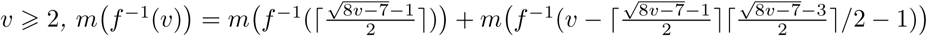.

*Proof.* The number of leaves in the tree of CP rank *v* ⩾ 2, or *m*(*f* ^−1^(*v*)), is the sum of the numbers of leaves in its left and right subtrees, or *m*(*k*_1_(*v*)) + *m*(*k*_2_(*v*)). ▪

The CP ranking, unlike the Furnas ranking, assigns trees whose numbers of leaves differ substantially to neighboring ranks (Table 3). Unlike the Furnas ranking, however, it enables a straightforward calculation of the rank associated with a given tree and the tree associated with a given rank.

## 3 Smallest CP rank for a fixed number of leaves

Next, we compute the CP ranks of the trees of size *n* that have the smallest and largest CP ranks. For *n* ⩾ 1, we define 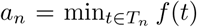 and 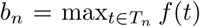. The sequences 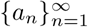 and 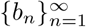 give the minimal and maximal CP rank considering all trees of size *n* leaves. Let *z*_*n*_ and *Z*_*n*_ respectively denote the trees of size *n* that achieve the minimal and maximal CP rank, *f* (*z*_*n*_) = *a*_*n*_ and *f* (*Z*_*n*_) = *b*_*n*_.

We begin with *a*_*n*_. To determine a recursion for *a*_*n*_, we first must establish that *a*_*n*_ increases with *n*.

### Lemma 4.

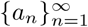 *is a strictly increasing sequence.*

*Proof.* First, by the definition of the CP ranking and the fact that tree sizes *n* = 1, 2, and 3 each have only one tree, *a*_1_ = 1, *a*_2_ = 2, and *a*_3_ = 3. We show by induction that for each *n* ⩾ 3, *a*_*n*+1_ ⩾ *a*_*n*_.

Consider a tree *t* of size *n* + 1. We must show *f* (*t*) > *a*_*n*_, as it would then follow that 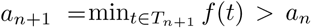. We consider two cases. (i) Suppose the two subtrees of the root of *t* have sizes *n* and 1. Then the left subtree of *t* has size *ℓ*(*t*) = *n* and the right subtree has size 1, and

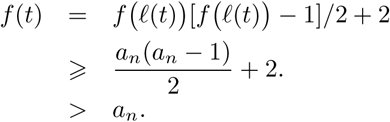

Here, the first inequality uses *f (ℓ*(*t*)) ⩾ *a*_*n*_ by the definition of *a*_*n*_, and the second follows from the quadratic inequality *x*(*x* − 1)*/*2 + 2 > *x*. Thus, each tree *t* of size *n* + 1 with subtrees of size *n* and 1 has *f* (*t*) > *a*_*n*_.

(ii) Suppose *t* instead has subtrees of size 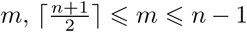, and *n* + 1 − *m* ⩽ *m*. The subtrees of *t* are *ℓ*(*t*) and *r*(*t*), one of which has size *m* and the other of which has size *n* + 1 − *m* (possibly *m* = *n* + 1 − *m* for odd *n*). As it is not yet specified which subtree is *ℓ*(*t*) and which is *r*(*t*), we consider both left–right arrangements, in each exhibiting a tree *t*′ of size *n* with *a*_*n*_ ⩽ *f* (*t*′) < *f* (*t*).

Suppose that *ℓ*(*t*) has size *m*. Then *f (ℓ*(*t*)) ⩾ *f* (*z*_*n*+1−*m*_) by the inductive assumption: if *ℓ*(*t*) has size *m*, then *f (ℓ*(*t*)) ⩾ *a*_*m*_⩾ *a*_*n*+1−*m*_. Consider a tree *t*′ of size *n* whose two subtrees are *ℓ*(*t*) and *z*_*n*−*m*_. Note that *f (ℓ*(*t*)) ⩾ *a*_*m*_ > *a*_*n*−*m*_ = *f z*_*n*−*m*_ by the inductive assumption, so that the canonical form for *t*′ has *ℓ*(*t*′) = *ℓ*(*t*′) and *r*(*t*′) = *z*_*n*−*m*_. We then have

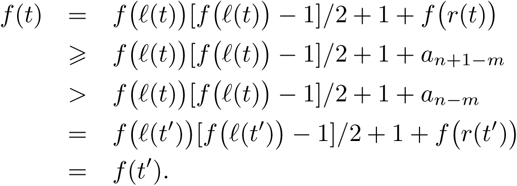

The first inequality follows from the definition of *a*_*n*_, and the second follows from the inductive assumption. Thus, *f* (*t*) > *f* (*t*′) *a*_*n*_.

Now suppose instead that *ℓ*(*t*) has size *n* + 1 − *m*. Let the two subtrees of *t*′ be *r*(*t*) and *z*_*n*−*m*_. Then *f* (*r*(*t*)) *a*_*m*_ > *a*_*n*−*m*_ = *f* (*z*_*n*−*m*_), so that in canonical form, *t*′ has *ℓ*(*t*′) = *r*(*t*) and *r*(*t*′) = *z*_*n*−*m*_. We have

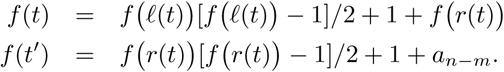

It follows that *f* (*t*) > *f* (*t*′) is equivalent to [*f (ℓ*(*t*)) − *f (r*(*t*))][*f (ℓ*(*t*)) + *f (r*(*t*)) − 3] > 2[*a*_*n*−*m*_ − *f (ℓ*(*t*))]. This latter inequality holds, as *f (ℓ*(*t*)) − *f (r*(*t*)) ⩾ 0 for any *t, f (ℓ*(*t*)) + *f (r*(*t*)) − 3 ⩾ 0 for any *t* with *m*(*t*) ⩾ 3, and *f (ℓ*(*t*)) ⩾ *a*_*n*+1−*m*_ > *a*_*n*−*m*_ by the inductive hypothesis. Thus, *f* (*t*) > *f* (*t*′) ⩾ *a*_*n*_.

We conclude that for each tree of size *n* + 1 with subtrees of size *m* and 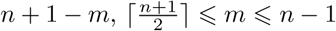, we can find a tree *t*′ of size *n* for which *f* (*t*) > *f* (*t*′). As *f* (*t*′) *a*_*n*_, it follows that *f* (*t*) > *a*_*n*_. ▪

The computation of *a*_*n*_ encodes a result that the tree with minimal CP rank is obtained by appending two subtrees of minimal CP rank for their size to a shared root. These subtrees are identical for even *n*, and they differ in size by one leaf for odd *n*.

### Theorem 5.

*The sequence* 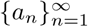 *of values of the minimal CP rank across trees of fixed size n satisfies*

a. *a*_1_ = 1.
b. *a*_2*n*_ = *a*_*n*_(*a*_*n*_ − 1)*/*2 + 1 + *a*_*n*_ *for* 2*n* ⩾ 2, *and*
c. *a*_2*n*−1_ = *a*_*n*_(*a*_*n*_ − 1)*/*2 + 1 + *a*_*n*−1_ *for* 2*n* − 1 ⩾ 3.

*Proof.* The base case of *a*_1_ = 1 is trivial, as are the cases of *a*_2_ = 2 and *a*_3_ = 3. Consider a tree *t* with an even number of leaves 2*n* ⩾ 4.

We claim that if *m (ℓ*(*t*)) < *n*, then *t* ≠ *z*_2*n*_. Suppose the left subtree of *t* has *n*^*^ < *n* leaves. The right subtree then has at least *n* + 1 leaves, so that *f (ℓ*(*t*)) > *f (r*(*t*)) ⩾ *a*_*n*+1_. Then *ℓ*(*t*) cannot equal 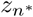, as 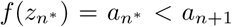 by Lemma 4. We could then construct a tree of 2*n* leaves whose left subtree is *r*(*t*) and whose right subtree is 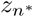. This tree would have a lower CP rank than *t*, as the inequality

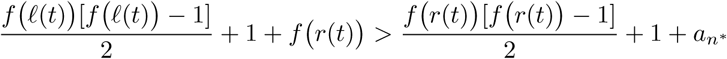

is equivalent to 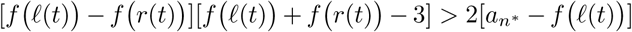; this latter inequality holds as its left side is nonnegative and its right size is negative. Thus, *m ℓ*(*z*_2*n*_) *n*.

Having established that the canonical form of *z*_2*n*_ has *m ℓ*(*z*_2*n*_) *n*, we have (*ℓ*(*z*_2*n*_)) ∈ *T*_*n*_ ∪ *T*_*n*+1_ ∪ … ∪ *T*_2*n*−1_. We now argue that *z*_2*n*_ is the tree *t*^*^ whose left subtree is *z*_*n*_ and whose right subtree is also *z*_*n*_.

For *t, t*′ in canonical form, *f (ℓ*(*t*)) < *f (ℓ*(*t*′)) implies *f* (*t*′) < *f* (*t*′) (Section 2); for *t, t*′ in canonical form with *f (ℓ*(*t*′)) = *f (ℓ*(*t*′)) and *f (r*(*t*)) < *f (r*(*t*′)), *f* (*t*) < *f* (*t*′). By Lemma 4, *a*_*n*_ ⩽ *a*_*n*+1_ ⩽ … ⩽ *a*_2*n*−1_, so that 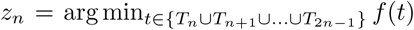. Combining these results, each tree *t* ≠ *t*^*^ with *t* ∈ *T*_2*n*_ and *ℓ*(*t*) ∈ *T*_*n*_ ∪*T*_*n*+1_ ∪…∪*T*_2*n*−1_, written in canonical form, has *f* (*t*) > *f* (*t*^*^): if *ℓ*(*t*) ≠ *z*_*n*_, then *f* (*t*) > *f* (*t*^*^); if *ℓ*(*t*) = *z*_*n*_ and *r*(*t*) ≠ *z*_*n*_, then *f* (*t*) > *f* (*t*^*^). We conclude *ℓ*(*z*_2*n*_) = *r*(*z*_2*n*_) = *z*_*n*_ and *a*_2*n*_ = *a*_*n*_(*a*_*n*_ − 1)*/*2 + 1 + *a*_*n*_.

For trees of size 2*n* − 1 ⩾ 5, the same argument applies: we show *m ℓ*(*t*) ⩾ *n*, then we argue that *z*_2*n*−1_ is the tree with left subtree *z*_*n*_ and right subtree *z*_*n*−1_, producing *a*_2*n*−1_ = *a*_*n*_(*a*_*n*_ − 1)*/*2 + 1 + *a*_*n*−1_. ▪

The first terms of 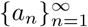 are 1, 2, 3, 4, 6, 7, 10, 11, 20, 22, 28, 29, 53, 56, 66, 67 (Table 2). The recursion for *a*_*n*_ constructs the trees *z*_*n*_. For odd *n*, the two subtrees immediately descended from the root of the tree *f* ^−1^(*a*_*n*_) have numbers of leaves that differ by 1 (Table 3). For even *n, f* ^−1^(*a*_*n*_) has two identical subtrees descended from the root. In both the odd and even cases, for each internal node, the two subtrees immediately descended from the node differ by at most 1 in their numbers of leaves. Note that in the case that *n* is a power of 2, *n* = 2^*k*^ for *k* ⩾ 1, the tree that has minimal CP rank is the fully symmetric tree.

## 4 Largest CP rank for a fixed number of leaves

We now turn to 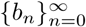, the sequence of values of the maximal CP rank among trees with *n* leaves. As in Section 3, we begin by demonstrating that *b*_*n*_ increases with *n*.

### Lemma 6.

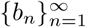 *is a strictly increasing sequence.*

*Proof.* We show *b*_*n*+1_ > *b*_*n*_ for *n* ⩾ 1.

For *n* ⩾ 1, we append *z*_*n*_ and *z*_1_ to a shared root to obtain a tree *t*. Then *f* (*t*) = *b*_*n*_(*b*_*n*_ − 1)*/*2 + 2. The inequality *b*_*n*_(*b*_*n*_ − 1)*/*2 + 2 > *b*_*n*_ always holds, as 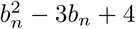 is an upward-facing parabola with vertex at a positive value, 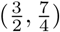. Thus, we have constructed a tree of *n* + 1 leaves with CP rank greater than that of the tree of *n* leaves with largest CP rank. ▪

Next, to obtain *b*_*n*_, we show that the tree of size *n* with maximal CP rank is obtained by appending the tree of maximal CP rank with size *n* − 1 and a single leaf to a shared root.

### Theorem 7.

*The sequence* 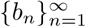 *of values of the maximal CP rank across trees of fixed size n satisfies*

a. *b*_1_ = 1.
b. *b*_*n*_ = *b*_*n*−1_(*b*_*n*−1_ − 1)*/*2 + 2 *for n* ⩾ 2.

*Proof.* The base case *b*_1_ = 1 is trivial, as are the cases of *b*_2_ = 2 and *b*_3_ = 3.

Let *n* ⩾ 4 and consider a tree *t* with *m*(*t*) = *n*. We claim that if 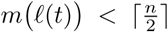, then *t* ≠ *Z*_*n*_. Suppose the left subtree of *t* has 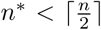 leaves. The right subtree then has at least 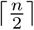 leaves, so that 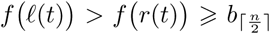. Then *ℓ*(*t*) cannot be 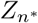, as 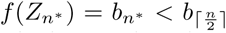 by Lemma 6. We could then construct a tree *t*′ of size *n* whose left subtree is 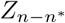 and whose right subtree is 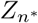. This tree would have a greater CP rank than *t*, as

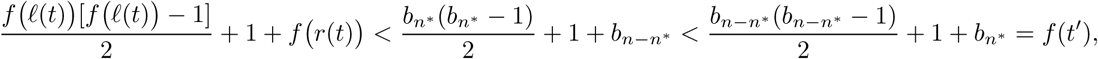

where we use 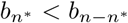 by Lemma 6. Thus, *m*(*ℓ*(*Z*_*n*_)) ⩾ *n*.

Having established that the canonical form of *Z*_*n*_ has *m*(*ℓ*(*Z*_*n*_)) ⩾ *n*, we have 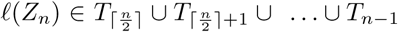. We now argue that *Z*_*n*_ is the tree *t*^*^ whose left subtree is *Z*_*n*−1_ and whose right subtree is *Z*_1_.

For *t, t*′ in canonical form, *f* (*ℓ*(*t*)) < *f* (*ℓ*(*t*′)) implies *f* (*t*) < *f* (*t*′) (Section 2). By Lemma 6, 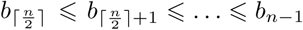, so that 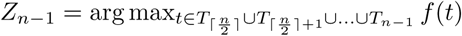.Combining these results, each tree *t* ≠ *t*^*^ with *t* ∈ *T*_*n*_ and 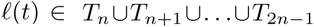, written in canonical form, has *f* (*t*) < *f* (*t*^*^). We conclude *ℓ*(*Z*_*n*_) = *Z*_*n*−1_ and *r*(*Z*_*n*_) necessarily is *Z*_1_. Hence *b*_*n*_ = *b*_*n*−1_(*b*_*n*−1_ − 1)*/*2 + 2. ▪

The first values of 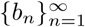 are 1, 2, 3, 5, 12, 68, 2280, 2598062 (Table 2, A108225 in OEIS). Because the tree *Z*_*n*_ with maximal CP rank is obtained by successively appending the tree *Z*_*n*−1_ with maximal CP rank and a single leaf to a shared root, the tree of *n* leaves that achieves maximal CP rank is the *caterpillar* tree—the tree in which there exists an internal node that descends from all other internal nodes (Table 3).

A relationship exists between entries of 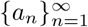 and entries of 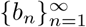. We write 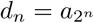 for *n* 0.

### Proposition 8.

*For n* ⩾ 0, *d*_*n*_ + 1 = *b*_*n*+2_.

*Proof.* We demonstrate the result by induction. We have *d*_0_ + 1 = *a*_1_ + 1 = 2 and *b*_2_ = 2. For the inductive step, we assume *d*_*n*_ + 1 = *b*_*n*+2_ and show *d*_*n*+1_ + 1 = *b*_*n*+3_.

By Theorem 5, for 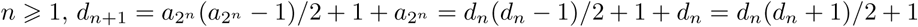. At the same time, *b*_*n*+3_ = *b*_*n*+2_(*b*_*n*+2_ − 1)*/*2 + 2 by Theorem 7. By the inductive hypothesis, we then have *b*_*n*+3_ = (*d*_*n*_ + 1)*d*_*n*_*/*2 + 2 = *d*_*n*+1_ + 1.

The sequence 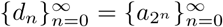 begins 1, 2, 4, 11, 67, 2279, 2598061 (A006894 in OEIS). As a result of Theorem 7 and Proposition 8, as we traverse ranks in the interval [*b*_*n*_, *b*_*n*+1_), flanked by the largest ranks for trees with *n* and *n* + 1 leaves, we encounter ranks for trees representing numbers of leaves as high as 2^*n*−1^. The CP ranking can place trees with quite different numbers of leaves in adjacent ranks. We characterize this difference in the following remark.

### Remark 9.

*For n* ⩾ 1, *all trees with CP rank in* [*b*_*n*_, *b*_*n*+1_) *have sizes in* [*n*, 2^*n*−1^]. *The smallest size for a tree with CP rank in* [*b*_*n*_, *b*_*n*+1_) *is n, and the largest size for a tree with CP rank in* [*b*_*n*_, *b*_*n*+1_) *is* 2^*n*−1^.

*Proof.* The interval [*b*_*n*_, *b*_*n*+1_), ranging from the largest CP rank of a tree with *n* leaves to one less than the largest CP rank of a tree with *n* + 1 leaves, contains the smallest CP rank of a tree with 2^*n*−1^ leaves (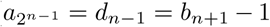 by Proposition 8). Because 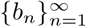 is increasing by Lemma 6, *b*_*n*−1_ < *b*_*n*_, so that no trees of size *n* − 1 leaves or fewer have CP rank in [*b*_*n*_, *b*_*n*+1_). Because 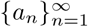 is increasing by Lemma 4, 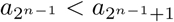, and no trees of size 2^*n*−1^ + 1 or greater have CP rank in [*b*_*n*_, *b*_*n*+1_). ▪

## 5 Asymptotics

We now evaluate the asymptotic behavior of the sequences 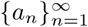 and 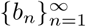. We use the method of Aho and Sloane [1].

### Theorem 10.

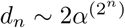 *for a constant α* ≈ 1.24602.

*Proof.* In Proposition 8, 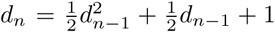 for *n* ⩾ 1, with *d*_0_ = 1. Substituting 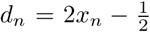, we obtain 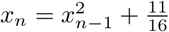, with 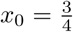.

We take *y*_*n*_ = log *x*_*n*_ in this quadratic recursion for *x*_*n*_. We then have, for *n* ⩾ 1, *y*_*n*_ = 2*y*_*n*−1_ + *α*_*n*−1_, where 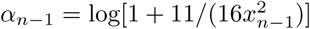. Applying the method of Aho and Sloane [1] for quadratic recursions,

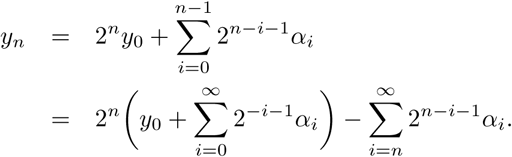

Exponentiating both sides, we obtain

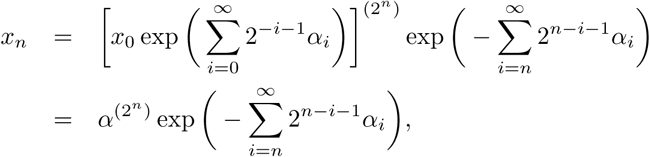

where *α* is the constant 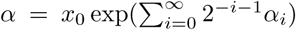. Inserting the first terms of the recursive sequence 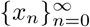, we have 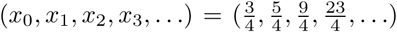. From these values, we have 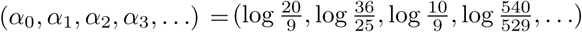. Numerically evaluating the constant *α* from the first 10 terms, we obtain *α* ≈ 1.24602083298366.

Then

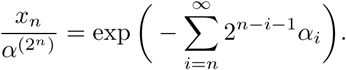

As *n* → ∞, the sum 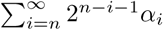 can be bounded 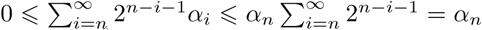. Because *x*_*n*_ → ∞ as *n* → ∞, *α*_*n*_ → 0 as *n* → ∞. Hence 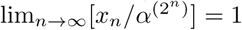.

Because 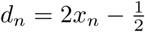, we conclude 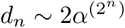. ▪

The connection between 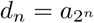 and *b*_*n*_ (Proposition 8) quickly gives the following result.

### Corollary 11.

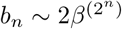 *for a constant β* ≈ 1.05653.

*Proof.* By Proposition 8, *b*_*n*_ ∼ *d*_*n*−2_, and by Theorem 10, 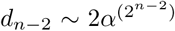. Hence, 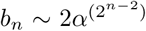. Writing *β* = *α*^1*/*4^ ≈ 1.05652876566960, the result follows. ▪

We have obtained an asymptotic equivalence for 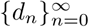 in Theorem 10, giving the increase of 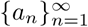 for the subsequence *n* = 1, 2, 4, 8, 16, We now place a bound on the increase in 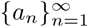 more generally.

### Proposition 12.

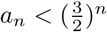 *for n* ⩾ 1.

*Proof.* We use induction. The result holds for 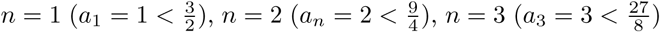, and 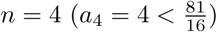. We assume that the inequality holds for each *n* from 1 to 2*k* − 2.

For even 2*k* ⩾ 4, applying Theorem 5 and the inductive hypothesis,

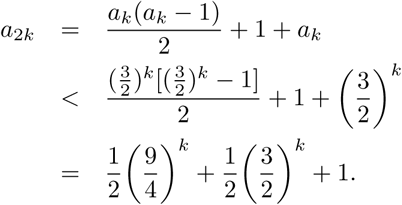

To demonstrate 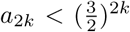, we must show 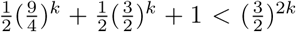, or equivalently, 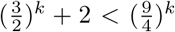 This latter inequality holds: 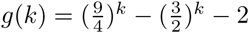 is an increasing function for *k* > 0, with 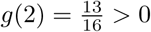, and *g*(*k*) therefore remains positive for *k* ⩾ 2.

For odd 2*k* − 1 ⩾ 5, applying Theorem 5 and the inductive hypothesis,

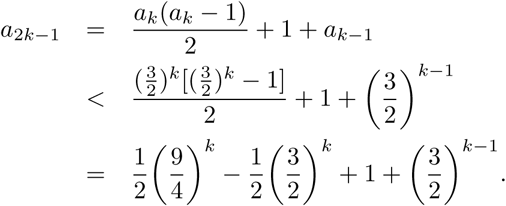

To demonstrate 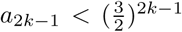, we must show 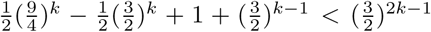, or equivalently, 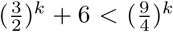. Again, the function 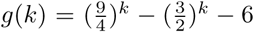 is increasing for *k* > 0, with 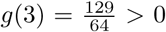. Hence, *g*(*k*) remains positive for *k* ⩾ 3.

In Figures 1 and 2, we examine the ratios *a*_*n*_*/*(2*α*^*n*^) and 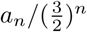 for small values of *n*. In Figure 1, the ratio *a*_*n*_*/*(2*α*^*n*^), which has limit 1 for the subsequence *n* = 1, 2, 4, 8, 16, … (Theorem 10), generally exceeds 1, returning to near 1 when *n* is equal to a power of 2. In Figure 2, the ratio 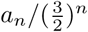 lies substantially below 1, indicating that 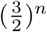 is a relatively loose upper bound for *a*_*n*_.

**Figure 1.**
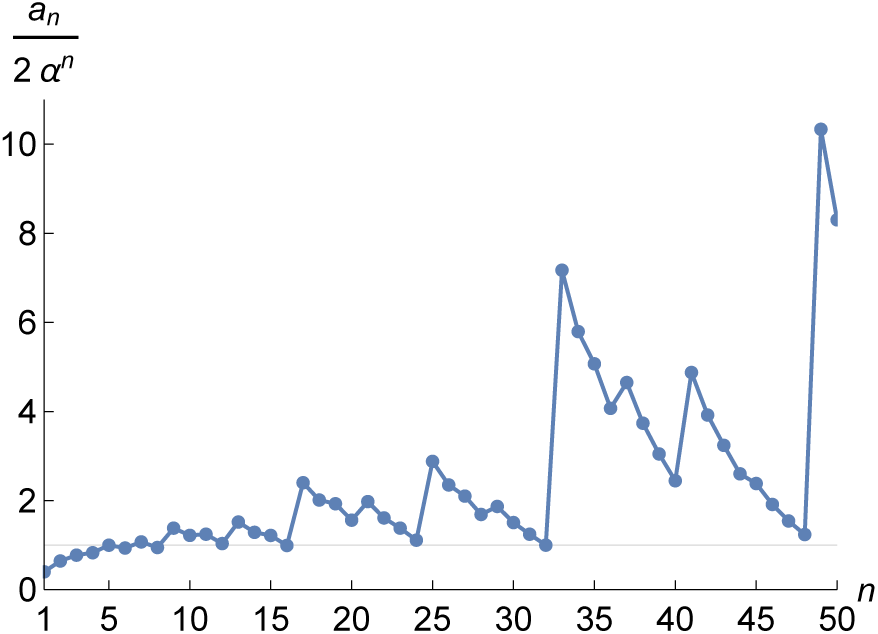
The ratio *a*_*n*_*/*(2*α*^*n*^), *α* ≈ 1.24602. This ratio has limit 1 for subsequence 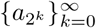 (Theorem 10).

**Figure 2.**
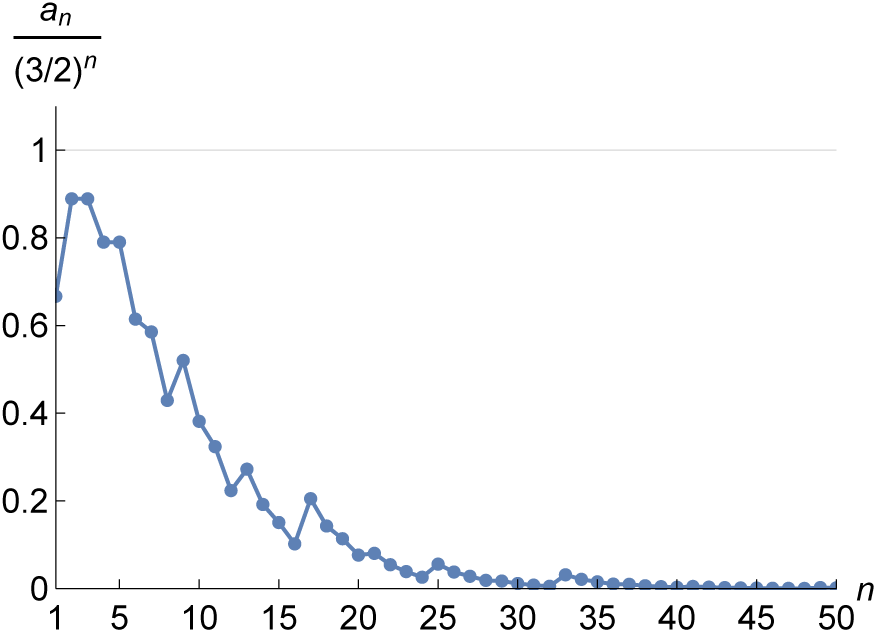
The ratio 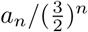. This ratio lies below 1 for all *n* (Proposition 12).

## 6 Discussion

The Colijn-Plazzotta ranking provides a convenient method for obtaining the rank associated with a given tree and the tree associated with a given rank. We have obtained recursions for the minimal and maximal CP rank across trees with *n* leaves (Theorems 5 and 7), analyzing their asymptotic behavior (Section 5). This analysis demonstrates that as the CP rank increases, the numbers of leaves in the associated trees traverse a wide range of values. In fact, for *n* ⩾ 1, the interval bounded by the largest rank across trees with *n* leaves and the largest rank across trees with *n* + 1 leaves contains ranks for trees with as many as 2^*n*−1^ leaves (Remark 9). Unlike for the Furnas ranking, the CP ranking has the property that the trees associated with sequential ranks do not necessarily differ in size by either 1 or 0 leaves; the difference in size between trees with sequential ranks is 2^*n*^ − *n* − 2 in the transition from rank 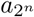 to rank 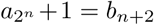. Asymptotically, the largest rank across trees with *n* leaves increases with 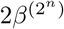 for a constant *β* ≈ 1.05653 (Corollary 11), and the smallest rank across trees with *n* leaves is bounded above by the substantially smaller 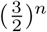 (Proposition 12), with asymptotic equivalence to 2*α*^*n*^, *α* ≈ 1.24602, for the subsequence 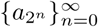 (Theorem 10).

The computations of *a*_*n*_ and *b*_*n*_ construct the trees *z*_*n*_ and *Z*_*n*_ that respectively have the smallest and largest CP ranks among *n*-leaf trees. The largest rank belongs to the caterpillar. The smallest rank belongs to a “balanced” tree, in which, for each internal node, the two subtrees descended from the node have either equally many leaves, or numbers of leaves that differ by 1. Thus, because the most extreme CP ranks among trees of size *n* are represented by a balanced tree and the unbalanced caterpillar tree, CP rank has potential to be useful in the measurement of tree balance—the extent to which an unlabeled shape resembles balanced shapes [2, 10, 12, 15]. Because the tree of minimal CP rank has absolute difference 0 or 1 between the sizes of the two subtrees for each internal node, it is perhaps useful to consider CP rank specifically in relation to the Colless tree balance index [3, 5, 6, 14]—which for each node sums the absolute difference in the numbers of descendants of the two subtrees of the node and which has larger values for unbalanced trees.

The study augments recent results examining unlabeled binary rooted trees that possess maximal or minimal features in scenarios arising from consideration of evolutionary problems [6, 7, 8, 13]. Curiously, Theorem 10 has a close connection with an analysis of “non-equivalent ancestral configurations,” structures that are used in characterizing relationships of pairs of trees [9, 16]. For non-equivalent ancestral configurations associated with the completely balanced trees—the same trees that produce the smallest CP rank in the case that *n* is a power of 2—Section 4.2 of Disanto & Rosenberg [9] gives a recursion for a quantity *γ*_*n*_, with *γ*_0_ = 0, which when transformed by 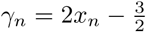 produces the recursion 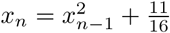 with 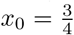 seen in the proof of Theorem 10. Thus, Disanto & Rosenberg [9] obtain the same asymptotic result 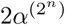 we observed, but for the growth of a different quantity, the number of non-equivalent ancestral configurations with increasing numbers of leaves 2^*n*^ in completely balanced trees.

The CP ranking encodes an innovative scheme that facilitates computations with unlabeled binary rooted trees, as shown by Colijn & Plazzotta [4] in their construction of metrics for unlabeled binary rooted trees and their use of these metrics to study evolutionary trees of strains of infectious agents. Further analysis of the mathematical properties of the CP ranking can potentially inform its applications.

## Acknowledgments

Support was provided by NIH grant R01 GM131404.

## References

[1] A. V. Aho and N. J. A. Sloane. Some doubly exponential sequences. Fibonacci Q., 11:429–437, 1973.

[2] M. G. B. Blum and O. François. On statistical tests of phylogenetic tree imbalance: the Sackin and other indices revisited. Math. Biosci., 195:141–153, 2005.

[3] G. Cardona, A. Mir, and F. Rosselló. Exact formulas for the variance of several balance indices under the Yule model. J. Math. Biol., 67:1833–1846, 2013.

[4] C. Colijn and G. Plazzotta. A metric on phylogenetic tree shapes. Syst. Biol., 67:113–126, 2018.

[5] D. H. Colless. Phylogenetics, the theory and practice of phylogenetic systematics. Syst. Zool., 31:100–104, 1982.

[6] T. M. Coronado, M. Fischer, L. Herbst, F. Rosselló, and K. Wicke. On the minimium value of the Colless index and the bifurcating trees that achieve it. q-bio.PE:1907.05064v2, 2020.

[7] F. Disanto and N. A. Rosenberg. Enumeration of ancestral configurations for matching gene trees and species trees. J. Comput. Biol., 24:831–850, 2017.

[8] F. Disanto and N. A. Rosenberg. Enumeration of compact coalescent histories for matching gene trees and species trees. J. Math. Biol., 78:155–188, 2019.

[9] F. Disanto and N. A. Rosenberg. On the number of non-equivalent ancestral configuriations for matching gene trees and species trees. Bull. Math. Biol., 81:384–407, 2019.

[10] J. Felsenstein. Inferring Phylogenies. Sinauer, Sunderland, MA, 2004.

[11] G. W. Furnas. The generation of random, binary unordered trees. J. Classif., 1:187–233, 1984.

[12] S. B. Heard. Patterns in tree balance among cladistic, phenetic, and randomly generated phylogenetic trees. Evolution, 46:1818–1826, 1992.

[13] A. Mir, F. Rosselló, and L. Rotger. A new balance index for phylogenetic trees. Math. Biosci., 241:125–136, 2013.

[14] A. Mir, L. Rotger, and F. Rosselló. Sound Colless-like balance indices for multifurcating trees. PLoS One, 13:e0203401, 2018.

[15] M. Steel. Phylogeny: Discrete and Random Processes in Evolution. Society for Industrial and Applied Mathematics, Philadelphia, 2016.

[16] Y. Wu. Coalescent-based species tree inference from gene tree topologies under incomplete lineage sorting by maximum likelihood. Evolution, 66:763–775, 2012.

